# Type 2B VWD substitutions trigger enhanced macrophage-mediated clearance that is attenuated by the BT200 aptamer

**DOI:** 10.64898/2026.09.07.749810

**Authors:** Alain Chion, Ciara Byrne, Anne-Marije Hulshof, Timon Albrecht, Bogdan Baci, Patricia Lopes, Ellie Karampini, Lidia Puertas-Umbert, Thomas A.J. McKinnon, Jamie M. O’Sullivan, Roger J.S. Preston, Shuhao Zhu, James C. Gilbert, Bernd Jilma, Ross I. Baker, Ferdows Atiq, James S. O’Donnell

## Abstract

Type 2B von Willebrand disease (VWD) is characterized by missense variants in exon 28 of the von Willebrand factor (VWF) gene that encodes the VWF-A1 domain. These missense mutations result in single amino acid residue substitutions that cause activation of VWF-A1 and promote spontaneous interaction with platelet GPIbα. Importantly, enhanced VWF clearance has been shown to play a key pathogenic role in >90% patients with Type 2B VWD. Although this VWF clearance occurs via mechanisms that are independent of VWF-platelet complex formation, the biological mechanisms responsible for the increased clearance of type 2B VWD variants remain poorly understood. In this study, we investigated a series of different type 2B amino acid substitutions within the VWF-A1 domain (R1306W, R1308C, W1313C and R1379L). We demonstrate that these variants all exhibit significantly increased macrophage binding. In part, the enhanced macrophage interactions are mediated via increased binding of type 2B VWD variants to LRP1 extracellular cluster II and IV. Our findings further demonstrate that the K1405-K1408 lysine cluster in the VWF-A1 domain plays a key role in enabling enhanced LRP1 interactions for type 2B VWD variants. In addition, we show that type 2B VWF variants demonstrate significantly enhanced interaction with the macrophage MGL receptor. Finally, and importantly from a clinical perspective, we demonstrate that the increase in macrophage binding for type 2B variants is significantly attenuated in the presence of BT200. Collectively, our findings have direct translational relevance with respect to the clinical heterogeneity and treatment of type 2B VWD.

## INTRODUCTION

Enhanced von Willebrand factor (VWF) clearance has been shown to play a key pathogenic role in >40% of patients with type 1 von Willebrand disease (VWD).^1–3^ Consequently, recent guidance from the ISTH Scientific and Standardization Subcommittee on VWF recommended that type 1 VWD patients with enhanced VWF clearance should be sub-classified into a distinct type 1C (1-Clearance) group.^4^ In addition, previous studies have shown that significantly increased VWF clearance is also common in patients with type 2 VWD.^5–8^ Indeed, the Willebrand in the Netherlands (WiN) study reported that the VWFpp/VWF:Ag ratio was actually higher in patients with type 2 compared to type 1 VWD.^5^ This finding suggests that enhanced clearance may play an even greater pathogenic role in type 2 VWD. Overall, >50 different amino acid substitutions in the *VWF* gene have been associated with increased clearance in VWD patients.^2,8^ Critically however, the molecular and cellular mechanisms through which these single amino acid substitutions cause enhanced clearance of multimeric VWF remain poorly understood.

Accumulating data have highlighted a key role for the VWF-A1 domain in regulating macrophage-mediated VWF clearance in vivo. For example, VWF binding to macrophages was significantly enhanced in the presence of either shear stress or ristocetin, suggesting a role for the VWF-A1 domain.^9–12^ Moreover, plate-binding assays have demonstrated that the VWF-A1 domain can bind directly to several clearance receptors including the low-density lipoprotein receptor-related protein-1 (LRP1),^10,13,14^ scavenger receptor class A member I (SR-A1)^15^ and macrophage galactose-type lectin (MGL).^16,17^ Recently, we demonstrated that a conserved cluster of four lysine residues (K1405, K1406, K1407 and K1408) within the VWF-A1 domain constitute a critical binding site for macrophage LRP1.^18^ Rondaptivon pegol (previously BT200) is a PEGylated RNA aptamer that interacts with the VWF-A1 domain and significantly attenuates VWF clearance in vivo.^19,20^ Furthermore, clinical studies have reported that BT200 can significantly increase plasma VWF and FVIII levels in healthy human controls and in patients with mild hemophilia A or VWD.^20–22^ Importantly, we have recently shown that BT200 interacts with the VWF-A1 domain in close proximity to K1405-K1408 lysine cluster and thereby attenuates macrophage LRP-1 mediated clearance.^18^ Finally, consistent with the concept that the VWF-A1 domain plays an important role in regulating VWF clearance, many of the VWF amino acid substitutions associated with type 1C VWD are clustered around the VWF-A1 domain.^2^

Type 2B VWD is characterized by missense variants clustered within exon 28 of the *VWF* gene that encodes the VWF-A1 domain.^23–26^ Single residue substitutions associated with type 2B VWD induce activation of VWF-A1 and promote spontaneous interaction with platelet GPIbα.^23,25,27,28^ Interestingly, clinical data have shown that many type 2B VWD amino acid substitutions are also associated with enhanced VWF clearance in vivo.^5,7,29^ For example, significantly enhanced VWF clearance was reported in 95% of type 2B VWD patients in WiN study.^5^ Furthermore, Wohner *et al* also demonstrated that the type 2B VWD variants VWF-R1306Q and VWF-V1316M were associated with significantly enhanced binding to LRP1.^29^ In this study, we investigated the biological mechanisms responsible for the increased clearance of type 2B VWD variants. We demonstrate that a series of different type 2B amino acid substitutions within the VWF-A1 domain directly result in enhanced macrophage binding. In part, these macrophage interactions are mediated via increased binding of type 2B VWD variants to LRP1 extracellular cluster II and IV respectively. Our findings further demonstrate that the K1405-K1408 lysine cluster in the VWF-A1 domain plays a key role in enabling enhanced LRP1 interactions for type 2B VWD variants. In addition, we show that type 2B VWF variants also demonstrate significantly enhanced interaction with the macrophage MGL receptor. Finally, and importantly from a clinical perspective, we demonstrate that the increase in macrophage binding for type 2B variants is significantly attenuated in the presence of BT200.

## MATERIALS AND METHODS

### Expression and purification of type 2B VWD variants

Recombinant wild type human VWF with a C-terminal His-tag was expressed in HEK293T cells using the expression vector pcDNA-VWF as previously described.^11,18^ Site-directed mutagenesis of human VWF was performed to introduce four specific type 2B amino acid substitutions (R1306W, R1308C, W1313C and R1379L) in the VWF-A1 domain. Subsequently, the lysine residues K1405, K1406, K1407 and K1408 in these type 2B variants were replaced with alanine residues to create VWF-R1306W-4A, VWF-R1308C-4A, VWF-W1313C-4A and VWF-R1379L-4A. Mutagenesis studies were performed using the Quickchange method with KOD Hot Start DNA polymerase (Merck, Cork, Ireland) and all mutations were confirmed by DNA sequencing. All VWF variants were transiently expressed in HEK293T cells. For each variant, conditioned serum free medium was harvested, concentrated and purified via nickel affinity chromatography, as described before.^13,30^

### Expression of LRP1 and MGL in HEK293 cells

Full-length LRP1 cDNA (13.63-Kb) was amplified on human placental cDNA (Invitrogen, Thermo Fisher Scientific) with KOD-Hot Start DNA Polymerase (Merck). PCR product was cloned into pJET vector (Thermo Fisher Scientific). LRP1 cDNA was excised by XhoI-PmeI digestion and ligated into pcDNA3.1-V5-HisA (Thermo Fisher Scientific). The LRP1 expression vector was then transfected into HEK293 with TansIT© 293 reagent (Mirus Bio) and clonal selection was performed using G418. MGL was transiently transfected in HEK293T using Turbofect (ThermoFisher Scientific) following manufacturer’s instructions. Briefly, 5 µg of MGL plasmid construct was added to 250 µl of Opti-MEM (gibco, ThermoFisher scientific) followed by addition of 10 µl transfection reagent. After 20 min incubation at room temperature, the solution is added to a T75 flask containing HEK293T in fresh MEM-α supplemented with 10% FBS. Following 48h incubation, transfected cells were collected for flow cytometry experiments.

### VWF variant binding to macrophages, HEK-LRP1 and HEK-MGL

All murine experiments were approved by the Royal College of Surgeons in Ireland Ethics Committee (REC1315) and were performed in full accordance with the Health Product Regulatory Authority, Ireland (license AE19136/P060 and P081). In brief, bone marrow was isolated from femurs and tibias of male and female C57BL/6J mice (20-25g) and differentiated into macrophages following culturing in 10% macrophage colony-stimulating factor derived from L929-producing cells, fetal bovine serum (10%), and penicillin/streptomycin (1%) supplemented RPMI (Merck). Macrophage colony-stimulating media was supplemented on days 3 and 7 and bone marrow-derived macrophages (BMDMs) were collected on day 10 for further experiments. BMDMs were resuspended at 200,000 cells per condition and incubated with VWF variants at 37°C for 30 minutes, followed by murine Fc Block (Miltenyi biotec). Live cells were identified using far-red fluorescent viability dye. Cell-bound VWF was detected using rabbit polyclonal anti-VWF DAKO antibody, followed by Alexa Fluor 488-conjugated goat anti-rabbit IgG secondary antibody. HEK-LRP1 and -MGL cells were detached using Accutase (10%) and VWF binding was assessed as described above, without Fc blocking. For experiments involving BT200 aptamer, VWF variants (10 µg/mL) were pre-incubated with BT200 (0-3000 nM) for 30 minutes at 37°C before addition to the cells. Fluorescence intensity was measured on a BD FACSCanto (BD Biosciences) or NxT Attune flow cytometer (Bio-Science, ThermoFisher Scientific). Mean fluorescent intensity (MFI) was analyzed using FlowJo software.

### VWF variant binding to LRP1 cluster II, cluster IV and MGL

For immunosorbent plate-binding assays, LRP1 cluster II was coated at 2μg/ml, LRP1 cluster IV was coated at 1 μg/ml and MGL was coated at 5μg/ml. Plates were blocked using 3% BSA and 1% polyvinylpyrrolidone (PVPP) solution in TBS-T with 2.5 mM CaCl_2_. Next, full-length VWF variants were incubated for 2 hours at 37°C in TBS-T containing 1% BSA, 0.33% PVPP with 2.5mmol/L CaC_l2_ (incubation buffer). VWF binding was subsequently detected using rabbit polyclonal anti-VWF DAKO-HRP conjugated antibody diluted 1/2000 in incubation buffer.

### Type 2B variant clearance studies in VWF^-/-^ mice

*VWF^-/-^* mice on a C57Bl/6J background were obtained from the Jackson Laboratory (Sacramento, USA). *VWF^-/-^* mice were infused with 7.5 µg of full-length recombinant human VWF or 2B variants via tail-vein injection. Blood was collected via cardiac puncture and samples were acquired following injection (time = 0min), and specific time points to generate an in vivo clearance curve. Residual VWF:antigen (VWF:Ag) levels were determined as before. The results of the in vivo clearance studies are presented as percentage residual VWF:Ag levels over time.

### Data Presentation and Statistical Analysis

Plate- and cell-binding curves were plotted using “nonlinear regression curve fit”. Data points represent mean values and error bars illustrate standard error of the mean (SEM). The extra-sum-of-squares F test was used to statistically compare regression curves. A P value <0.05 was considered statistically significant. All graphs were visualized using GraphPad Prism 10.2.0 (GraphPad Software LLC) and PyMOL (Schrödinger LLC) was used to generate a structural model of the VWF-A1 Domain visualizing the spatial localization of the K1404-1408 cluster and evaluated 2B variants.

## RESULTS

### VWF binding to macrophages is significantly enhanced in type 2B VWD

Previous clinical studies in patients using VWFpp/VWF:Ag ratios and fall-off after DDAVP have reported that accelerated VWF clearance is common in patients with type 2B VWD.^5,7^ To validate the hypothesis that specific type 2B VWD variants are associated with enhanced clearance, we first expressed two type 2B variants VWF-R1306W and VWF-R1308C respectively. In vivo studies performed in *VWF^-/-^* mice following tail vein injection confirmed that the clearance of VWF-R1306W and VWF-R1308C were both significantly increased compared to wild type VWF (**Figures 1A-1B**). Interestingly, the clearance of VWF-R1308C was also significantly (p=0.0063) faster compared to VWF-R1306W.

**Figure 1.**
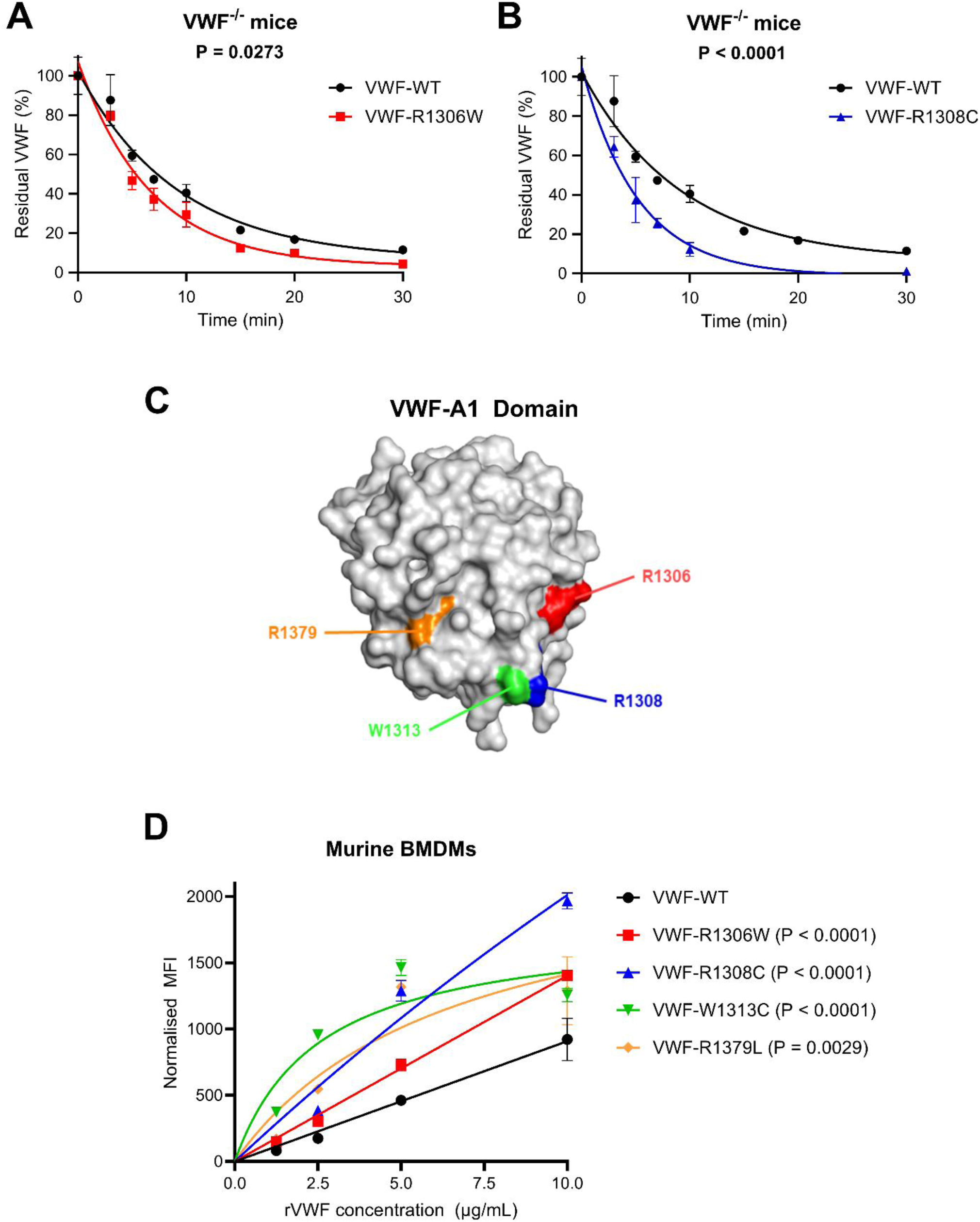
VWD-2B variants have increased in vivo clearance and macrophage binding. In vivo clearance of recombinant von Willebrand factor (rVWF) wild-type (WT) compared to the von Willebrand disease (VWD)-2B variants **(A)** R1306W and **(B)** R1308C following tail vein injection in VWF^-/-^ mice. Presented is the residual plasma VWF concentration at set post-infusion intervals. Three to seven mice per timepoint were used. **(C)** PyMOL model highlighting the spatial localization of R1306, R1308, W1313 and R1379 within the VWF-A1 domain. **(D)** Comparison VWF-WT murine bone-marrow derived macrophage (BMDM) binding compared to the VWD-2B variants R1306W, R1308C, W1313C, and R1379L using flow cytometry. BMDM binding is expressed as normalized mean fluorescent intensity (MFI) across different recombinant VWF concentrations. P-values illustrate the extra-sum-of-squares F test of VWF-2B variants compared to VWF-WT.

Hepatic macrophages play a critical role in regulating VWF clearance in vivo.^9,11,15^ To assess the role of macrophages in mediating VWF clearance in type 2B VWD, we next expressed four common type 2B variants (VWF-R1306W, VWF-R1308C, VWF-W1313C and VWF-R1379L) which are all located within the VWF-A1 domain (**Figure 1C)**. Consistent with previous studies,^9,13^ we confirmed using flow cytometry that wild type human VWF bound to murine bone-marrow derived macrophages (BMDM) in a concentration-dependent manner (**Figure 1D)**. Although there was some heterogeneity in macrophage interaction, significantly (p<0.005) enhanced BMDM binding was observed for each of the type 2B variants VWF-R1306W, VWF-R1308C, VWF-W1313C and VWF-R1379L (**Figure 1D)**. Consistent with our in vivo clearance data, binding of VWF-R1308C to murine BMDMs was also significantly (p=0.0005) enhanced compared to VWF-R1306W. Collectively, these data confirm that type 2B VWD amino acid substitutions within the VWF-A1 domain are associated with enhanced clearance in vivo and demonstrate enhanced macrophage-binding.

### LRP1 cluster II and cluster IV interact with type 2B VWD variants

VWF can interact with several different receptors on macrophages.^2^ In previous studies of two type 2B variants (VWF-R1306Q and VWF-V1316M), Wohner *et al* reported significantly enhanced binding to LRP1.^29^ To further investigate the role of LRP1 in regulating increased VWF clearance in type 2B VWD, full-length human LRP1 was stably expressed on HEK293 cells (HEK-LRP1) (**Figure 2A**). Recombinant wild type VWF bound to HEK-LRP1 cells in a concentration-dependent manner (**Figure 2B**). In keeping with our BMDM-binding results, heterogeneity was again seen between the different type 2B variants with respect to their ability to interact with HEK-LRP1 cells. Importantly however, significantly (p<0.0001) enhanced binding to HEK-LRP1 cells was observed for all four of the type 2B variants VWF-R1306W, VWF-R1308C, W1313C and R1379L studied (**Figure 2B)**.

**Figure 2.**
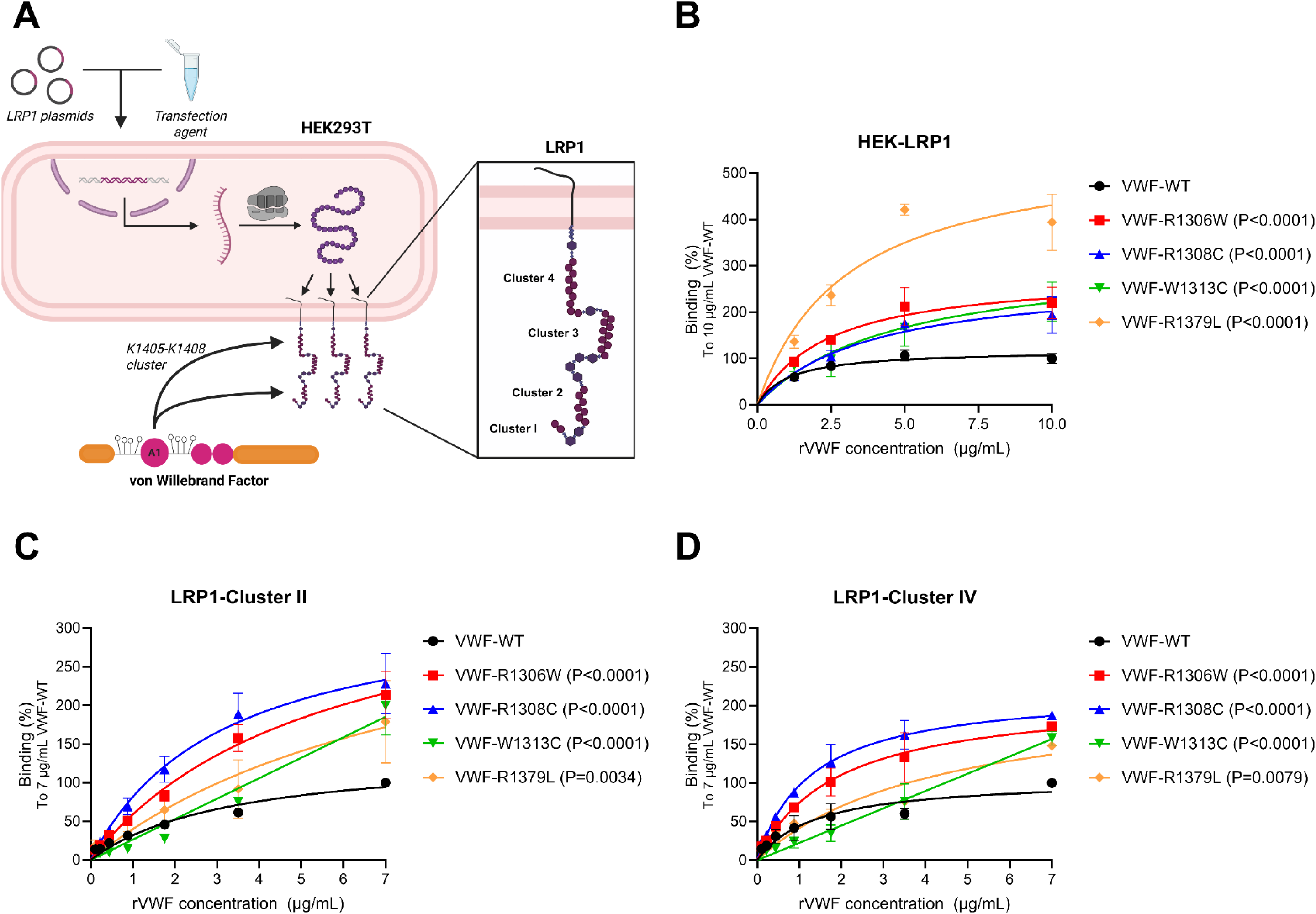
VWD-2B variants demonstrate increased LRP1 binding. **(A)** Overview of HEK293 transfection to induce membrane LRP1 receptor expression for in vitro binding experiments. The VWF-A1 domain K1405-K1408 lysine cluster binds LRP1 through cluster II & IV. **(B)** Comparison of von Willebrand factor (VWF) wild-type (WT) to HEK-LRP1 versus von Willebrand disease (VWD) 2B variants R1306W (red), R1308C (blue), W1313C (green), and R1379L (orange) using flow cytometry. HEK-LRP1 binding is expressed as percentage (%) mean fluorescent intensity compared to 10 µg/mL VWF-WT. Comparison of **(C)** LRP1-Cluster II and **(D)** LRP1-Cluster IV plate-binding between WT-VWF and 2B variants. Plate-binding across experiments was expressed as percentage (%) 450nm absorbance compared to 7 µg/mL VWF-WT. P-values illustrate the extra-sum-of-squares F test of VWF-2B variants versus VWF-WT.

The LRP1 receptor consists of four extracellular cluster repeats (I through IV).^31^ Recombinant wild type VWF bound to purified LRP1 cluster II and IV in plate-binding assays in a concentration-dependent manner (**Figures 2C-2D)**. Consistent with the HEK-LRP1 cellular binding studies, in plate-binding assays we observed that interactions of VWF-R1306W, VWF-R1308C, W1313C and R1379L to LRP1 cluster II were all significantly increased compared to wild type VWF (**Figure 2C**). Similarly, although the effect was less marked compared to cluster II, significantly enhanced binding of VWF-R1306W, VWF-R1308C, VWF-R1379L and VWF-W1313C to LRP1 cluster IV was also seen (**Figure 2D).** Together, these findings support the hypothesis that LRP1 contributes to macrophage-mediated enhanced VWF clearance in type 2B VWD and suggest that the LRP1 extracellular cluster II may play a specific role in this context.

### The K1405-K1408 lysine cluster in the VWF-A1 domain influences interaction of type 2B variants with LRP1

Recently, we demonstrated that the K1405-K1408 lysine cluster on the surface of the VWF-A1 domain constitutes a critical binding site in VWF for macrophage LRP1-mediated clearance (**Figures 3A-3B**).^18^ Thus, alanine mutagenesis of this lysine cluster (VWF-4A) significantly attenuated VWF binding to LRP1 and prolonged in vivo half-life.^18^ To assess whether the K1405-K1408 cluster is important for LRP1 interaction with type 2B VWD variants, we next studied the effect of removal of the K1405-K1408 cluster by alanine mutagenesis for each of the type 2B VWD variants VWF-R1306W-4A, VWF-R1308C-4A, VWF-W1313C-4A and VWF-R1379L-4A (**Figures 3A-3B**). Importantly, the enhanced binding of the type 2B variants to BMDMs was significantly inhibited following removal of the K1405-K1408 cluster (**Figures 3C-3F**). Similarly, enhanced binding to HEK-LRP1 cells was also abrogated following removal of the K1405-K1408 cluster (**Figure 3G**). Cumulatively, these data show that the K1405-K1408 lysine cluster in the VWF-A1 domain plays an important role in regulating enhanced LRP-1 mediated interactions in type 2B VWD.

**Figure 3.**
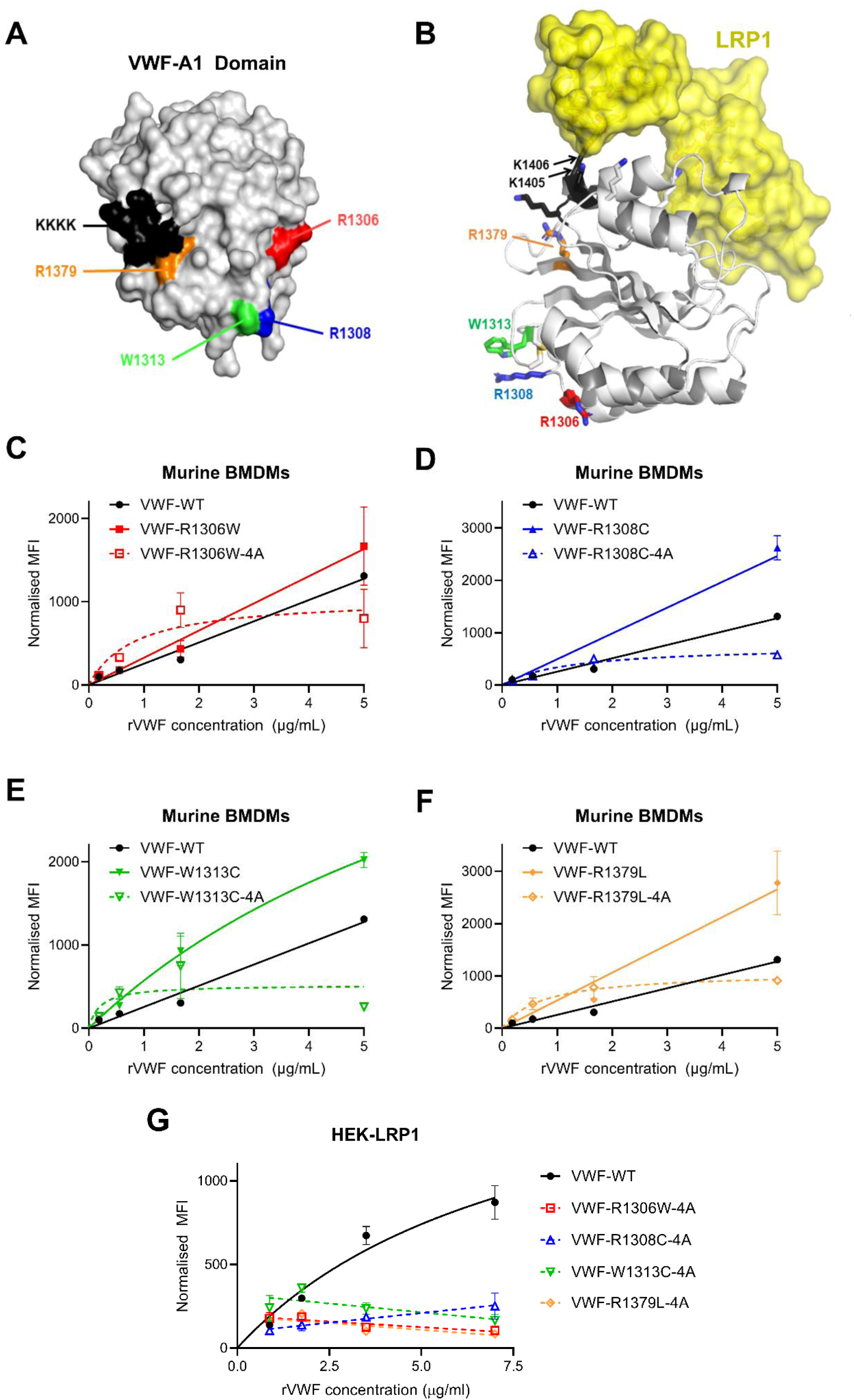
The K1405-K1408 cluster attenuates VWD-2B variant binding to BMDMs and HEK-LRP1. **(A)** PyMOL model illustrating the spatial localization of the K1405-K1408 cluster (black), R1306 (red), R1308 (blue), W1313 (green), and R1379 (orange) in the VWF-A1 domain. **(B)** Proposed interaction between VWF-A1 domain K1405-K1408 cluster and macrophage LRP1 receptor. Comparison of VWF-WT murine bone-marrow derived macrophage (BMDM) binding compared to the 2B variants using flow cytometry. BMDM binding is expressed as normalised mean fluorescent intensity (MFI) across different recombinant VWF (rVWF) concentrations. The enhanced binding of the von Willebrand disease (VWD) 2B variants **(C)** R1306W, **(D)** R1308C, **(E)** W1313C, and **(F)** R1379L to BMDMs was significantly inhibited following removal of the K1405-K1408 cluster (-4A variants). **(G)** Comparison of VWF-WT binding to HEK-LRP1 versus VWD-2B variants using flow cytometry. HEK-LRP1 binding is expressed as normalized MFI and removal of the K1405-K1408 cluster abolished HEK-LRP1 binding.

### BT200 attenuates interaction of type 2B variants with LRP1

Clinical studies have reported that the BT200 PEGylated RNA aptamer increases plasma VWF levels in healthy controls and in patients with mild haemophilia A.^21,32^ Furthermore, we recently showed that BT200 binds the VWF-A1 domain close to K1405-K1408 cluster and thereby inhibits macrophage LRP1-mediated clearance in vivo (**Figure 4A**).^18^ Importantly however, although BT200 has also been shown to increase plasma VWF levels in patients with type 2B VWD,^22^ the mechanisms underpinning this effect have not been defined. In keeping with previous data, we first confirmed that binding of wild type VWF to HEK-LRP1 cells was significantly inhibited by BT200 in a concentration-dependent manner (**Figure 4B**).^18^ Although heterogeneity was observed between different type 2B variants, nevertheless the enhanced binding of VWF-R1306W, VWF-R1308C, VWF-W1313C and VWF-R1379L to HEK-LRP1 cells were all significantly attenuated in the presence of BT200 (**Figures 4B-4E**). Importantly however, even in the presence of 3000nM BT200, the type 2B variants VWF-R1306W, VWF-R1308C, VWF-W1313C still demonstrated significantly increased residual HEK-LRP1 binding relative to wild type VWF. Altogether, these findings suggest that the increase in plasma VWF levels seen in type 2B patients treated with BT200 is likely due at least in part to reduced macrophage LRP1-mediated clearance.

**Figure 4.**
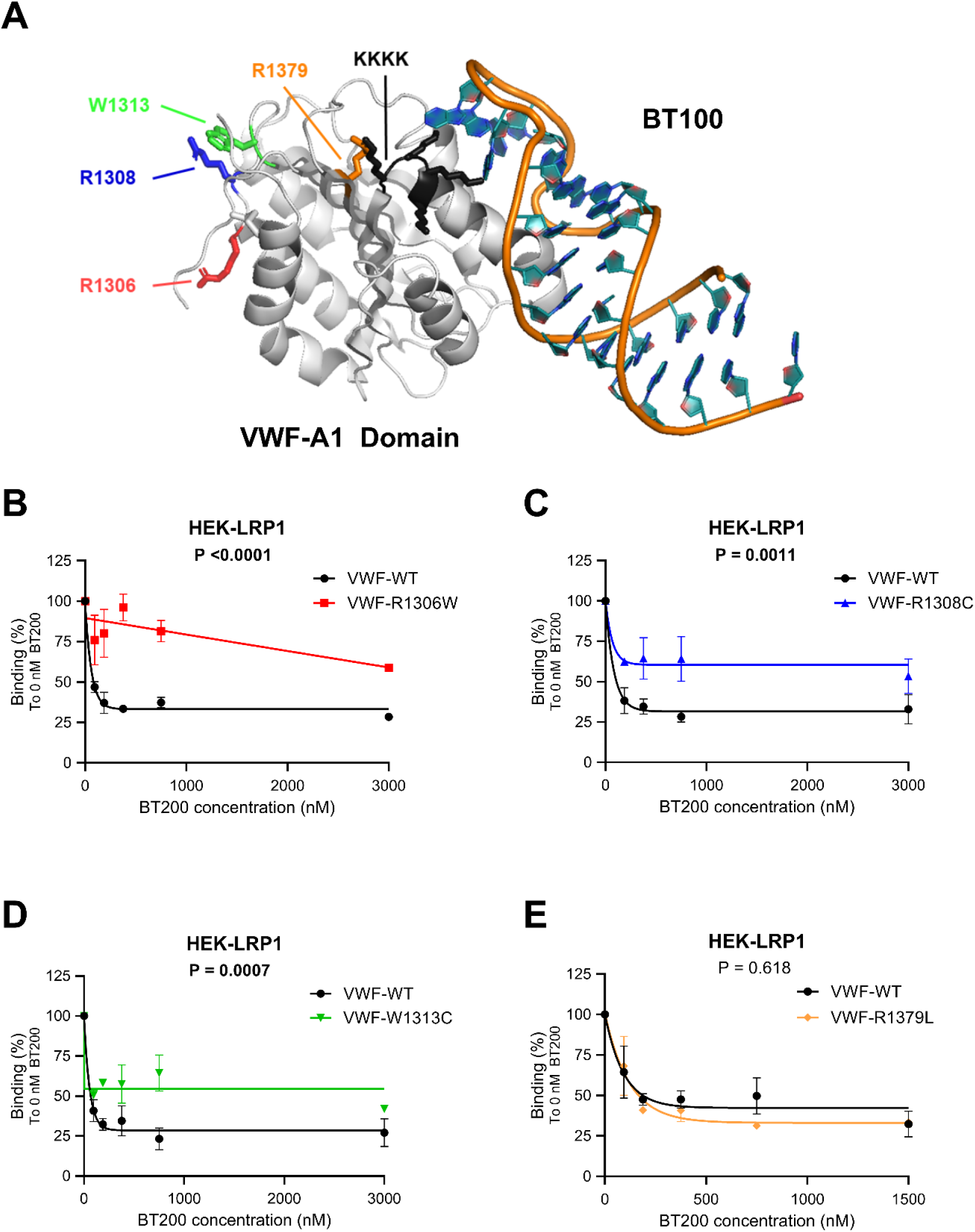
The aptamer BT200 attenuates VWF-2B variant binding to HEK-LRP1. **(A)** PyMOL model of VWF-A1 Domain binding to the aptamer BT100 (unpegylated BT200), highlighting the spatial localization of the K1405-K1408 cluster (black), R1306 (red), R1308 (blue), W1313 (green), and R1379 (orange). Comparison of VWF-WT HEK-LRP1 binding versus **(B)** R1306W, **(C)** R1308C, **(D)** W1313C, and **(E)** R1379L in the presence of the BT200 (0-3000nM) using flow cytometry. Residual recombinant VWF (rVWF) binding to HEK-LRP1 across different BT200 concentrations was calculated from the mean fluorescent intensity. Results are presented as residual HEK-LRP1 binding (%) compared to 10µg/mL rVWF. P-values illustrate the extra-sum-of-squares F test of VWF-2B variants compared to VWF-WT.

### The MGL clearance receptor contributes to enhanced clearance in type 2B VWD

Recent studies have shown that the MGL receptor on macrophages plays a role in regulating physiological VWF clearance and highlighted that the O-linked glycan that flank the VWF-A1 domain modulate MGL-mediated clearance **(Figure 5A)**.^16,17,33^ Furthermore, VWF binding to MGL was significantly enhanced in the presence of ristocetin.^16^ Given the effects of type 2B variants on VWF-A1 domain conformation, we hypothesized that MGL may also contribute to macrophage-mediated clearance in type 2B VWD. To investigate this hypothesis, human MGL was expressed on HEK293T cells (HEK-MGL) (**Figure 5A**). In keeping with previous studies, concentration-dependent binding of recombinant wild type VWF to HEK-MGL cells was observed (**Figure 5B**).^16,17^ Importantly, significantly (p<0.002) enhanced binding to HEK-MGL cells was observed for the type 2B variants VWF-R1306W, VWF-R1308C, VWF-W1313C and VWF-R1379L (**Figure 5B**). Consistent with the HEK-MGL cellular binding studies, interactions of VWF-R1306W, VWF-R1308C, VWF-W1313C and VWF-R1379L with purified MGL in immunosorbent plate-binding assays were also all significantly (p<0.0001) increased compared to wild type VWF (**Figure 5C**).

**Figure 5.**
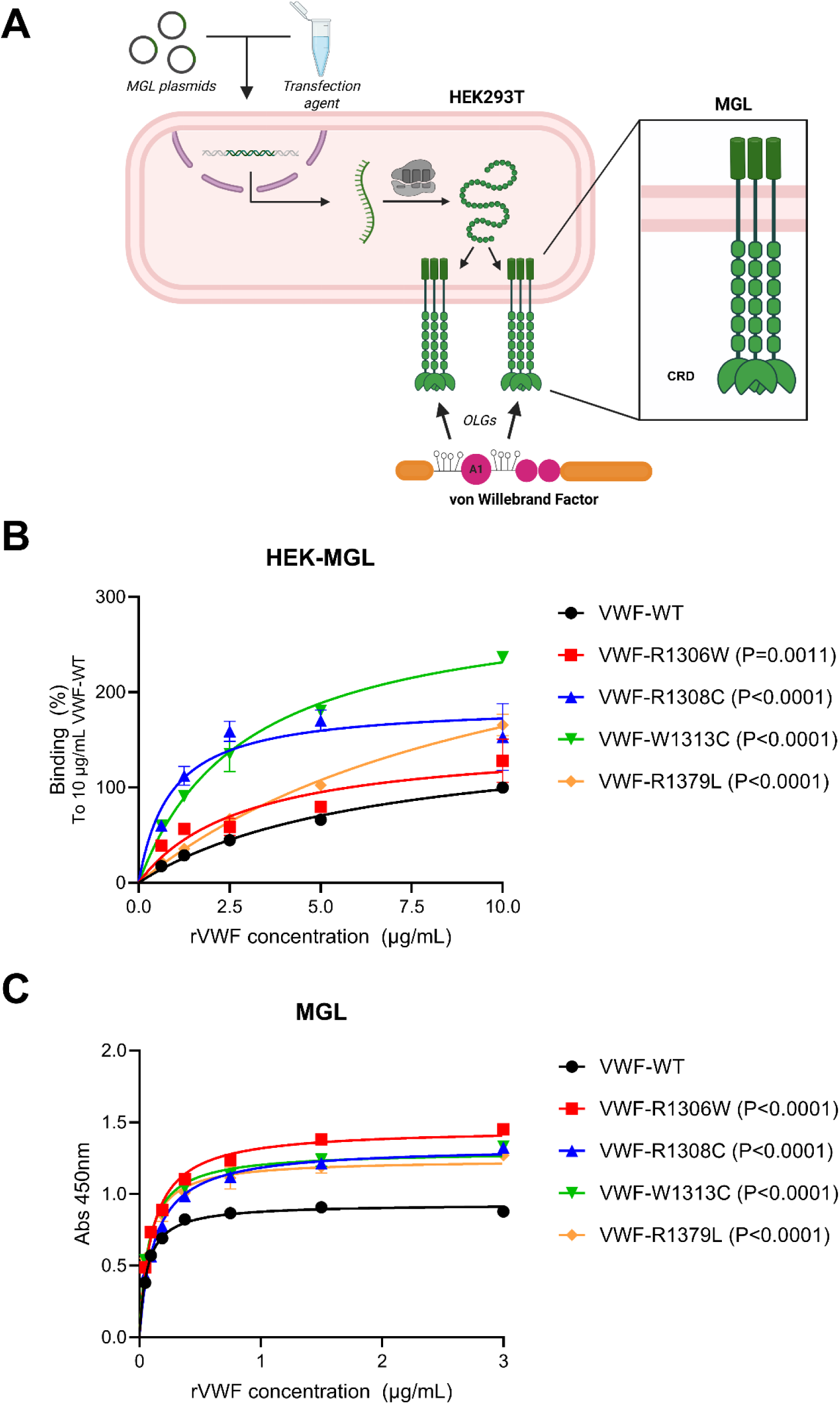
VWF-2B variants demonstrate increased MGL binding. **(A)** Overview of HEK293T transfection to induce membrane MGL receptor expression for in vitro binding experiments. The O-linked glycans (OLGs) flanking the VWF-A1 Domain mediate MGL binding. **(B)** Comparison of VWF wild-type (WT) binding to HEK-MGL versus von Willebrand disease (VWD) 2B variants R1306W (red), R1308C (blue), W1313C (green), and R1379L (orange) using flow cytometry. HEK-MGL binding across experiments was expressed as percentage (%) mean fluorescent intensity compared to 10 µg/mL VWF-WT. **(C)** Comparison of MGL plate-binding between WT-VWF and VWD-2B variants. Binding was presented as absorbance at 450nm across different recombinant VWF concentrations. P-values illustrate the extra-sum-of-squares F test of VWF-2B variants compared to VWF-WT.

In addition to inhibiting binding to LRP1, we recently showed that binding of pegylated BT200 to the VWF-A1 domain also attenuated the interaction of wild type human VWF with MGL.^18^ Moreover, this inhibitory effect was dependent upon the presence of the 40kD PEG moiety of BT200.^18^ Consequently, we next investigated whether pegylated BT200 influenced binding of type 2B variants to HEK-MGL cells. Although heterogeneity was again observed between the different type 2B variants, nonetheless the enhanced binding of VWF-R1306W, VWF-R1308C, VWF-W1313C and VWF-R1379L to HEK-MGL cells was significantly attenuated in the presence of BT200 (**Figures 6A-6D**). However, in keeping with the HEK-LRP1 data, we observed that the type 2B variant VWF-R1308C demonstrated significantly (p=0.0004) enhanced HEK-MGL binding relative to wild type VWF even in the presence of 300nM BT200. Together, these findings suggest that the increase in plasma VWF levels observed in type 2B patients treated with BT200 is mediated in part through a reduction in macrophage MGL-mediated macrophage clearance.

**Figure 6.**
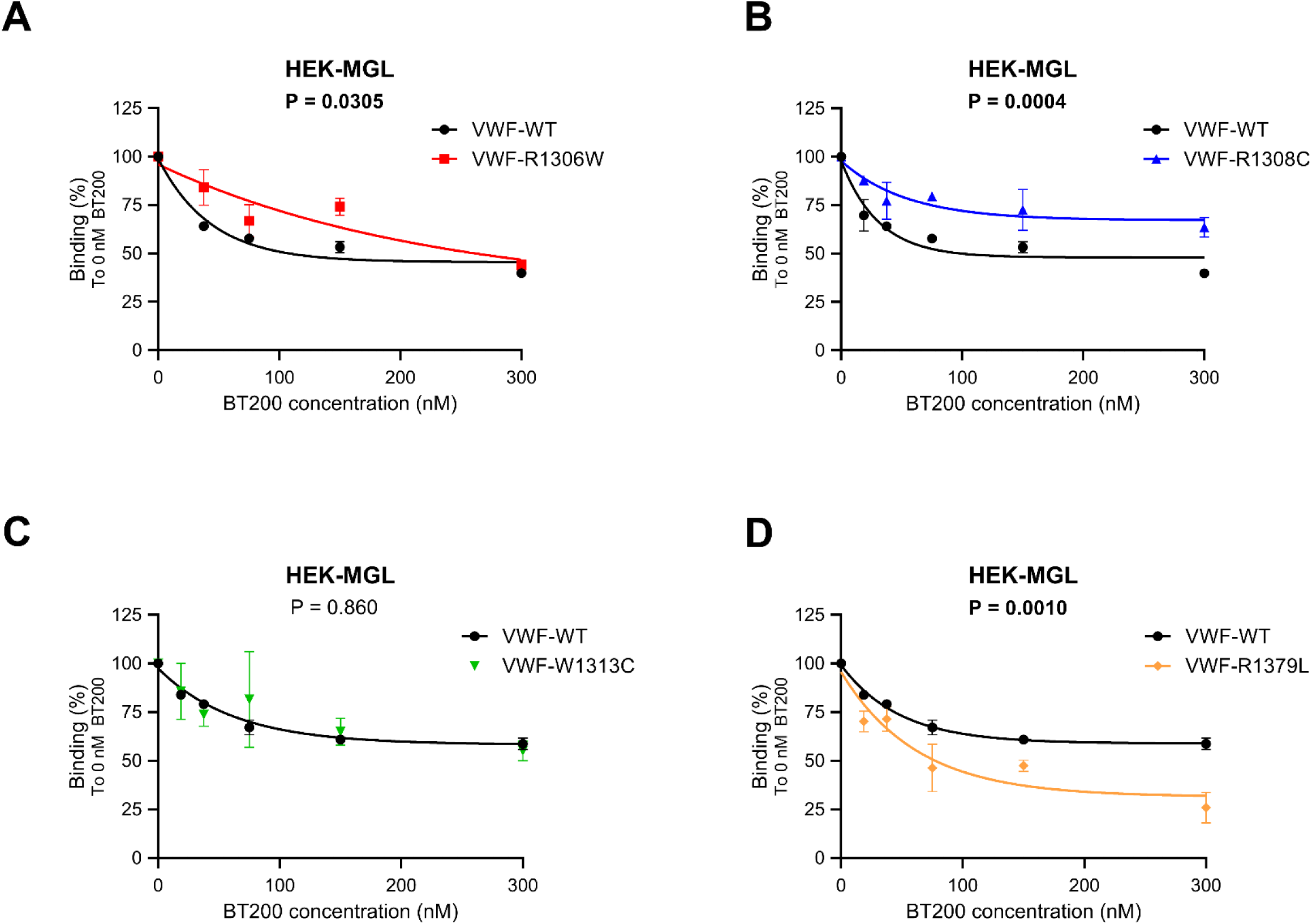
The aptamer BT200 attenuates VWF-2B variant binding to HEK-MGL. Comparison of VWF-WT HEK-MGL binding versus 2B variants **(A)** R1306W, **(B)** R1308C, **(C)** W1313C, and **(D)** R1379L using flow-cytometry. Residual recombinant VWF (rVWF) binding to HEK-MGL across different BT200 concentrations was calculated from the mean fluorescent intensity. Results are presented as residual HEK-MGL binding (%) compared to 10µg/mL rVWF. P-values illustrate the extra-sum-of-squares F test of VWF-2B variants compared to VWF-WT.

## DISCUSSION

Although increased clearance constitutes a key pathogenic mechanism in type 2B VWD,^5,7,8^ the underlying mechanisms involved have not been clearly defined. Casonato *et al* previously reported that VWF survival following DDAVP treatment was significantly reduced in patients with various type 2B variants compared to healthy controls.^7^ Furthermore, the reduced VWF half-life was independent of any changes in VWF multimer distribution or thrombocytopenia. Consistently, Wohner *et al.* subsequently showed that the half-lives of the type 2B variants VWF-R1306Q and VWF-V1316M in *VWF^-/-^* mice were both reduced compared to wild type VWF.^29^ Herein, we further demonstrate that clearance of the type 2B variants VWF-R1306W and VWF-R1308C is also significantly enhanced in *VWF^-/-^* mice. Importantly, these type 2B variants were expressed in human VWF which does not interact with murine GPIbα. Consequently, these observations support the hypothesis that enhanced clearance in type 2B VWD occurs, at least in part, via mechanisms that are independent of VWF-platelet complex formation.

Based on current evidence, it is clear that multiple single amino acid substitutions within the VWF-A1 domain induce conformational changes in type 2B VWD that result in gain-of-function interaction with platelet GPIbα.^1,23,25,27,34^ Critically however, these sequence variants and conformational changes in the VWF-A1 domain also trigger increased VWF clearance in vivo.^5,8^ Our findings provide novel insights into the biological mechanisms underpinning the enhanced clearance of type 2B VWD variants. In particular, we demonstrate that four different type 2B variants (VWF-R1306W, VWF-R1308C, VWF-W1313C and VWF-R1379L) demonstrate significantly enhanced binding to macrophages. With respect to specific macrophage cell surface receptors, we further show that enhanced interaction with LRP1 contributes to the increased clearance of type 2B VWD variants. This is mediated via increased binding of VWF to both LRP1 extracellular clusters II and cluster IV respectively. Furthermore, the K1405-K1408 region in the VWF-A1 domain plays a key role in enabling enhanced LRP-1 mediated clearance in type 2B VWD. Collectively, our data suggest that under steady-state conditions, the K1405-K1408 binding site for LRP1 in the VWF-A1 domain is not fully accessible in normal globular VWF. However, in the presence of shear or ristocetin, activation of the VWF-A1 domain facilitates not only enhanced GPIbα interaction but also promotes VWF clearance via LRP1. Similarly, in type 2B VWD, alterations in the VWF-A1 domain due to missense mutations enable spontaneous GPIbα binding and promote interaction of the K1405-K1408 cluster with LRP1 to drive increased macrophage-mediated clearance.

Besides enhanced LRP1-mediated clearance, our findings highlight that the MGL receptor also contributes to enhanced macrophage-mediated binding of type 2B VWD variants. Importantly, previous studies have shown that VWF binding to MGL is also significantly enhanced in the presence of ristocetin.^11,16,17^ Like both LRP1 and GPIbα, the MGL-binding site may therefore also be partly cryptic in multimeric VWF until the VWF-A1 domain is activated. In previous studies, we have demonstrated that sialylation of the O-linked glycans clustered at the N- and C-terminal ends of the VWF-A1 domain play a key role in regulating VWF interaction with MGL.^17^ In addition, Voos *et al* recently reported that the O-glycans on residues 1255 and 1256 are important for stability of the autoinhibitory module (AIM) that masks the A1 domain and regulates its binding to platelet GPIbα.^35^ Thus, we hypothesize that altered accessibility to O-linked glycan determinants due to AIM unfolding in type 2B VWD facilitates enhanced interaction with the MGL clearance receptor. Cumulatively, our findings therefore suggest that circulating multimeric VWF with a VWF-A1 domain that is in an active conformation for GPIbα interaction will also be targeted for rapid clearance by macrophages via (i) LRP-1 via enhanced K1405-K1408 exposure and (ii) MGL via altered O-linked glycans presentation (**Figure 7**). Given the potential for activated VWF-A1 domains to spontaneously bind to GPIbα and trigger platelet aggregate formation, the need for rapid clearance of activated VWF makes clear biological sense. In this context, the activated VWF-A1 domain is thus serving as an immunological damage-associated-molecular-pattern (DAMP) to drive macrophage-mediated clearance. Consistently, Casari *et al* previously reported that accelerated uptake of VWF-platelet complexes by macrophages was also an important regulator of thrombocytopenia in a murine model of type 2B VWD.^36^

**Figure 7.**
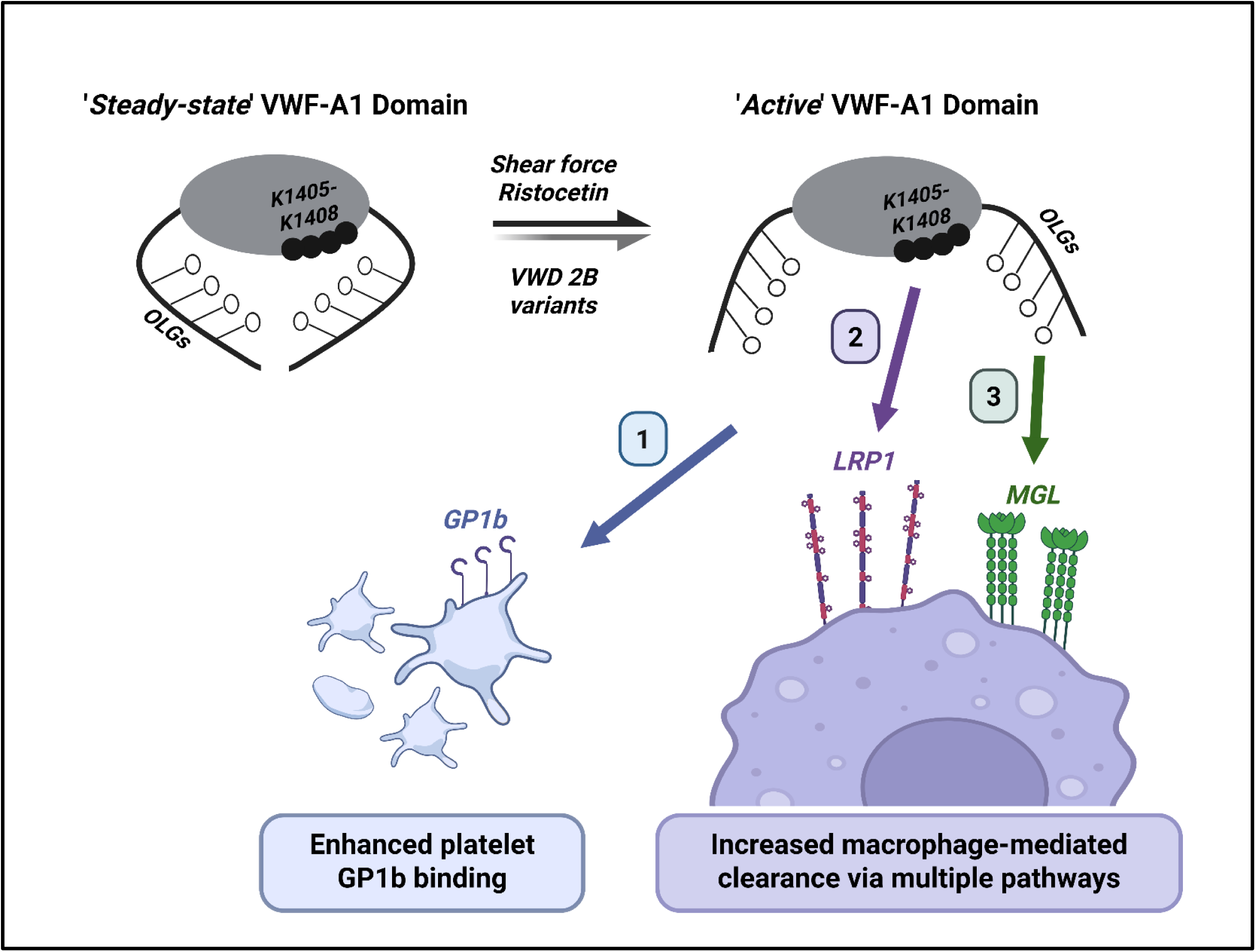
Altered accessibility VWF-A1 Domain mediates increased clearance in VWD type 2B. Our findings suggest that distinct von Willebrand disease (VWD) type 2B mutations induce conformational changes that alter the accessibility of the VWF-A1 domain lysine cluster K1406-K1408 and its flanking O-linked glycans (OLGs). This shift towards an ‘active’ VWF-A1 conformation promotes (1) increased platelet GPIb interaction, (2) enhanced macrophage-mediated clearance via LRP1 through enhanced K1405-1408 exposure, and (3) increased MGL-mediated clearance through altered presentation of OLGs.

Although all type 2B mutations are located within the VWF-A1 domain and cause enhanced binding to platelet GPIbα, nevertheless they are associated with significant differences in clinical phenotype, including differences in thrombocytopenia and/or relative reductions in HMWM.^5,26,34^ Herein, we demonstrate that different type 2B amino acid substitutions within the VWF-A1 domain are also associated with significant differences in macrophage-mediated clearance. For example, although the in vivo clearance of human VWF-R1306W and VWF-R1308C were both enhanced in *VWF^-/-^* mice, the clearance of VWF-R1308C was significantly faster. Consistently, binding of VWF-R1308C to murine BMDMs was also significantly (p=0.0005) enhanced compared to VWF-R1306W. Interestingly, Wohner *et al* previously also reported differences in clearance for two other type 2B variants (VWF-V1316M faster than VWF-1306Q).^29^ Our results further demonstrate that although type 2B variants demonstrate enhanced binding to both LRP1 and MGL clearance receptors, nonetheless there remains significant heterogeneity. However, again in keeping with our in vivo and BMDM binding data, we observed enhanced binding of VWF-R1308C to both HEK-LRP1 and HEK-MGL compared to VWF-R1306W. Together, these findings support the hypothesis that different type 2B variants cause differences in the VWF-A1 domain conformation that ultimately translate into variability in platelet GPIbα interaction, heterogeneity in macrophage clearance via LRP1- and MGL-mediated pathways, and ultimately differences in clinical phenotype. Finally, several other cell types including liver sinusoidal endothelial cells^37^ and hepatocytes^38^ have been shown to also play roles in regulating VWF clearance. Further studies will be required to determine whether these additional cell-mediated clearance pathways may also contribute to enhanced VWF clearance in type 2B VWD. Additional studies will also be needed to elucidate whether different type 2B variants influence interactions for other reported VWF-A1 binding ligands, or any of the emerging novel functions of VWF.^39^

Our findings have translational relevance with respect to the treatment of type 2B VWD. Previous studies have shown that the pegylated BT200 aptamer interacts with the VWF-A1 in proximity to the K1405-1408 cluster and thus attenuates macrophage-mediated clearance of wild-type VWF in healthy individuals.^18^ In addition, a recent phase 2 clinical trial demonstrated that BT200 significantly increased plasma VWF levels and platelet count in type 2B VWD patients.^22^ Of note, the type 2B patients in that study had the same VWF-A1 domain variants (VWF-R1306W, VWF-R1308C, and W1313C) that were compared in our study. Consistent with our data, significant heterogeneity in clinical responses was observed between the individual type 2B patients following BT200 treatment.^22^ These clinical data further support the hypothesis that different type 2B variants trigger enhanced macrophage-mediated clearance through different pathways. Further studies will be required to determine whether other VWF mutations clustered around the VWF-A1 domain in patients with type 1C or type 2A VWD may also trigger enhanced macrophage-mediated clearance.^2,8,40^ In addition, the efficacy of BT200 in attenuating pathological enhanced clearance of these different VWD variants will need to be compared to that of other emerging VWD therapies including the KB-V13A12 bispecific nanobody^41^ and the monovalent antibody HMB-002.^42^

## Acknowledgements

J.S.O’D is supported by a Science Foundation Ireland Frontiers for the Future (FFP) award (20/FFP-A/8952); by funds from the Royal City of Dublin Hospital Trust (Project grant 221) and by a Research Ireland Strategic Partnership Programme grant award (24/SPP/1340). F.A is supported by a Rubicon grant (452022310) from the Netherlands Organization for Health Research and Development (ZonMw). AMH is supported by funds of the EU Horizon Marie Skłodowska-Curie Actions (MSCA) Postdoctoral Fellowship (101202483). Cartoons were created with BioRender.com.

## Authorship contributions

AC, CB, TAJM, RJSP, SZ, JCG, BJ, RB, FA and JSOD designed the research; AC, CM, AMH, RJSP, JMOS, SZ, JCG, BJ, RB, FA and JSOD wrote the article; AC, AMH and FA performed statistical analysis; AC, CB, AMH, TA, BB, PL, EK, LPU and FA performed experiments, all authors contributed to final draft writing and critical revision. All authors participated sufficiently in this work, take public responsibility for the content, and approved the final version of the article.

